# *Urochloa trichopus* (Hoch.) Stapf (Poaceae: Paniceae): An addition to the Flora of Karnataka, India

**DOI:** 10.1101/2025.08.14.670282

**Authors:** Sandeep Pulla, Mukta Mande, Mahesh Sankaran

**Affiliations:** National Centre for Biological Sciences, Bengaluru, 560094, India

**Keywords:** Angiosperms, Gramineae, non-native, new record, Bengaluru

## Abstract

*Urochloa trichopus* (Hoch.) Stapf, a grass species native to Africa, the Arabian Peninsula, and Madagascar, is reported here for the first time from Karnataka (India). We provide a detailed taxonomic description with photographs and review its ecology.

## Introduction

*Urochloa* P.Beauv. (Family: Poaceae, Subfamily: Panicoideae, Tribe: Paniceae, Subtribe: Melinidinae) is a pantropical genus of over a hundred C4 grass species (Watson et al. 1992; Kellogg 2015). Of these, twenty-seven species and a further eight varieties are reported from India (Kellogg et al. 2020). The genus, *sensu lato*, is morphologically diverse, paraphyletic, and includes many species previously assigned to the genera *Brachiaria* (Trin.) Griseb., *Chaetium* Nees, *Scutachne* Hitchc. & Chase, *Megathyrsus* (Pilg.) B.K. Simon & S.W.L. Jacobs and *Eriochloa* Kunth (Kellogg 2015; Delfini et al. 2023; Masters et al. 2024). Many characters previously used to delimit the genus – mucronate upper lemma, abaxial spikelet orientation, transversely rugulose upper floret, racemose inflorescences – have subsequently been found to be of limited taxonomic value due to the occurrence of intermediate forms and high variability within the expanded genus (Bor 1960; Clayton and Renvoize 1986; Ashalatha and Nair 1993; Veldkamp 1996; Vorontsova 2022). As such, the genus is likely to undergo further revision (Vorontsova 2022). African *Urochloa* include several globally-important forage grasses such as *U. brizantha* (A.Rich.) R.D.Webster and *U. maxima* (Jacq.) R.D.Webster (≡ *Megathyrsus maximus* (Jacq.) B.K.Simon & S.W.L.Jacobs). Several African *Urochloa* are invasive in their introduced ranges and are known to negatively impact ecosystems (Pivello et al. 1999; Zenni and Ziller 2011; Rojas-Sandoval and Acevedo-Rodríguez 2022; Duque et al. 2022).

During routine grass collections for study, a small population (< 100 individuals) of *Urochloa* species was observed growing in an open-canopied, semi-wild habitat inside Gandhi Krishi Vigyan Kendra campus, Bengaluru (945 m ASL), amidst invasive plants such as *Parthenium hysterophorus* L. and *Calyptocarpus vialis* Less. and native plants such as *Setaria flavida* (Retz.) Veldkamp and *Sida spinosa* L. Examination of the literature indicated that the species was *U. trichopus* (Hochst.) Stapf, not previously reported from Karnataka (Sharma 1984; Karnataka Biodiversity Board 2019; Kellogg et al. 2020).

*U. trichopus* is a savanna woodland and grassland grass native to Africa, the Arabian Peninsula, and Madagascar. Morphologically and phylogenetically, it is closely allied to *U. mosambicensis* (Hack.) Dandy, which, together with *U. panicodies* P.Beauv. and *U. ramosa* (L.) T.Q.Nguyen form the Trichopus clade (Masters et al. 2024). Sosef (2016) noted that *U. trichopus* differs from *U. mosambicensis* largely by its annual, as opposed to perennial, habit. Deeming the distinction taxonomically insufficient, he synonymized the latter with the former. The Trichopus clade as a whole has significance as livestock feed.

*U. trichopus*’s ecology is not well-studied in its natural habitat. It appears to be prized by both wild and domesticated herbivores for its nutrient-rich tissues and is tolerant of intense herbivory (Van Oudtshoorn 1999; Treydte et al. 2013). A fast growing grass, it is considered an indicator of disturbed places and is associated with overgrazed and trampled areas and roadsides in southern Africa (Van Oudtshoorn 1999; Fish et al. 2015). The species grows well in light shade. In some regions, it dominates the herbaceous plant community after a fire (Heinl et al. 2007). A study in a southern African savanna found that, in contrast to other grass species, *U. trichopus* had greater mycorrhizal colonization in frequently burned, compared to unburned, areas (Hartnett et al. 2004). *U. trichopus* has been reported to be a highly-competitive invasive in Australia (Lawes and Grice 2010).

*U. trichopus* was introduced into India as a fodder crop (Bor 1960). It has been reported from Uttar Pradesh (Kellogg et al. 2020) and Telangana (Siddabathula and Yadav 2021) and, as *U. mosambicensis*, from Rajasthan and Tamil Nadu (Kabeer and Nair 2009; Kellogg et al. 2020). Given that many *Urochloa* fodder species are capable of being strongly competitive, producing allelopathic compounds, and altering fire regimes (Gorgone-Barbosa et al. 2015; Masters et al. 2024), *U. trichopus* merits careful evaluation of its potential as an invasive in India. Given the pronounced dormancy of *U. trichopus* seeds (McIvor and Howden 2000), their inadvertent transport with soil presents a plausible dispersal pathway.

## Methods

Specimens were processed by standard herbarium techniques and deposited at the National Centre for Biological Sciences. Micro-morphological characters were photographed using a Leica M125 C encoded stereo microscope (Leica Microsystems GmbH, Germany) and measured in Fiji (Schindelin et al. 2012).

### Taxonomic treatment

***Urochloa trichopus*** (Hochst.) Stapf in Oliv., Fl. Trop. Afr. 9: 589 (1920). ≡ *Panicum trichopus* Hochst. in Flora 27: 254 (1844). ≡ *Eriochloa trichopus* (Hochst.) Benth. in J. Linn. Soc., Bot. 19: 89 (1881). ≡ *Helopus trichopus* (Hochst.) Steud. in Syn. Pl. Glumac. 1: 100 (1854). = *Urochloa mosambicensis* (Hack.) Dandy in J. Bot. 69: 54 (1931). = *Panicum mosambicense* Hack., Bol. Soc. Brot. 6: 140 (1888). = *Echinochloa notabilis* (Hook. f.) Rhind, Grasses of Burma 50 (1945; as “notabile”). = *Panicum notabile* Hook. f., Fl. Brit. India (J.D. Hooker). 7(21): 32 (1896). = *Urochloa pullulans* Stapf, Fl. Trop. Afr. 9 (1920) 590, nom. superfl. = *Urochloa pullulans* Stapf var. *mosambicensis* Stapf, Fl. Trop. Afr. 9 (1920) 590, comb, incorr.

Caespitose perennials. Culms stoloniferous, geniculately ascending, 50—100 cm tall, nodes terete, bearded, lower nodes rooting; internodes 9.5—16.2 cm long. Leaves basal and cauline. Leaf sheath 6.1—13.0 × 0.2—1.3 cm, with longitudinal grooves, glabrous below, pubescent above, margins ciliate. Leaf blade 2.1—16.4 cm × 0.3—1.2 cm, chartaceous, lanceolate, base cordate to amplexicaul, apex acuminate, midrib prominent, margins tuberculate ciliate to glabrous, cilia ca. 2 mm. Ligule a fringe of hairs, 0.1—0.25 cm long. Inflorescence terminal or axillary, made up of racemes along a central axis, 7—35 cm long. Raceme axis 10.5—20.2 cm long, pubescent. Racemes 4—5(7), 3.3—6.2 × 0.2—0.4 cm, alternate, secund, spikelets imbricate, densely packed in two rows, disarticulating below glumes. Rachis 0.5 mm wide, wavy, pilose, margins winged, scabrid. Pedicel 0.3—0.7 mm long with 1-4 setae, 2.3—4.4 mm long. Spikelets 3.5—4.8 × 1.5—2.0 mm, abaxial, elliptic-ovate. Lower glumes 1.0—3.2 × 0.3—1.0 mm (2/3—3/4 length of spikelet), turned away from rachis, pubescent below, glabrous above, elliptic, apex rounded, 3-nerved, with 1—2 bristles on the mid-nerve inserted 2/3 from the base; bristles 1.0—2.0 mm long, geniculate, translucent white or purple. Upper glume 3.3— 4.3 × 1.4—1.9 mm, pubescent, elliptic-ovate, apex attenuate-acuminate, 3—5-nerved, 3—5-keeled. Florets 2; lower male, upper bisexual. Lower lemma 3.1—4 × 1.4—1.7 mm, hyaline, elliptic-ovate, apex acuminate, margins involute, fringed with purple bristles, 3—5-nerved, 3-keeled. Lower palea similar to lower lemma, 2.5—3.6 × 1.4—1.6 mm, hyaline, 2-nerved, without keels. Stamens 3, anthers 1.3—2.0 × 0.2—0.4 mm. Lodicules fringed, 0.2—0.3 mm long. Upper lemma 1.6—2.6 × 1.3—1.6 mm, elliptic, 3—5-nerved, without keels, transversely rugulose, apex mucronate, mucro 0.5—0.8 mm long, puberulent. Upper palea 1.5—2.8 × 1.2— 1.6 mm, hyaline, 3-nerved, apex obtuse, without keels, involute, winged, wings hyaline. Lodicules 0.2—0.3 mm long, cuneate, hyaline. Stamens 3. Ovary 0.5—1.2 × 0.2—0.9 mm. Stigmas 2, 1.2—1.8 × 0.2—0.5 mm, plumose, purple at maturity. Caryopsis 1.0—2.4 × 0.5— 1.5 mm, ovoid.

## Notes

*U. trichopus* is distinguished from *U. panicoides*, with which it shares abaxial spikelet orientation, by the lower glume being 2/3—3/4 the length of the spikelet (versus 1/4—1/3 in *U. panicoides*) and the lower lemma having a prominent fringe of bristles (versus not).

## Flowering & fruiting

June—October.

## India distribution

Rajasthan, Uttar Pradesh, Tamil Nadu, Telangana, Karnataka.

## Global distribution

Native to sub-Saharan Africa, Madagascar, Saudi Arabia, Yemen; introduced into India, Myanmar, Australia, Brazil, Argentina, Mexico, U.S.A.

## Specimens examined

India, Karnataka, Bengaluru, 13.08533 N 77.578289 E, 945 m ASL, 13-08-2025, *S. Pulla* 2 (JCB [HJCB 2000]) and 23-06-2025, *S. Pulla* 23VI25A (NCBS Research Collections [NRC-P-AA-0470], NCBS, Bengaluru).

## Acknowledgements

We are grateful to Dr. Navendu Page; Dr. Pritha Dey and Tarun Karmakar at NCBS Research Collections Facility; and Prof. Sankara Rao at Herbarium JCB for their support.

## Figures

**Figure 1.**
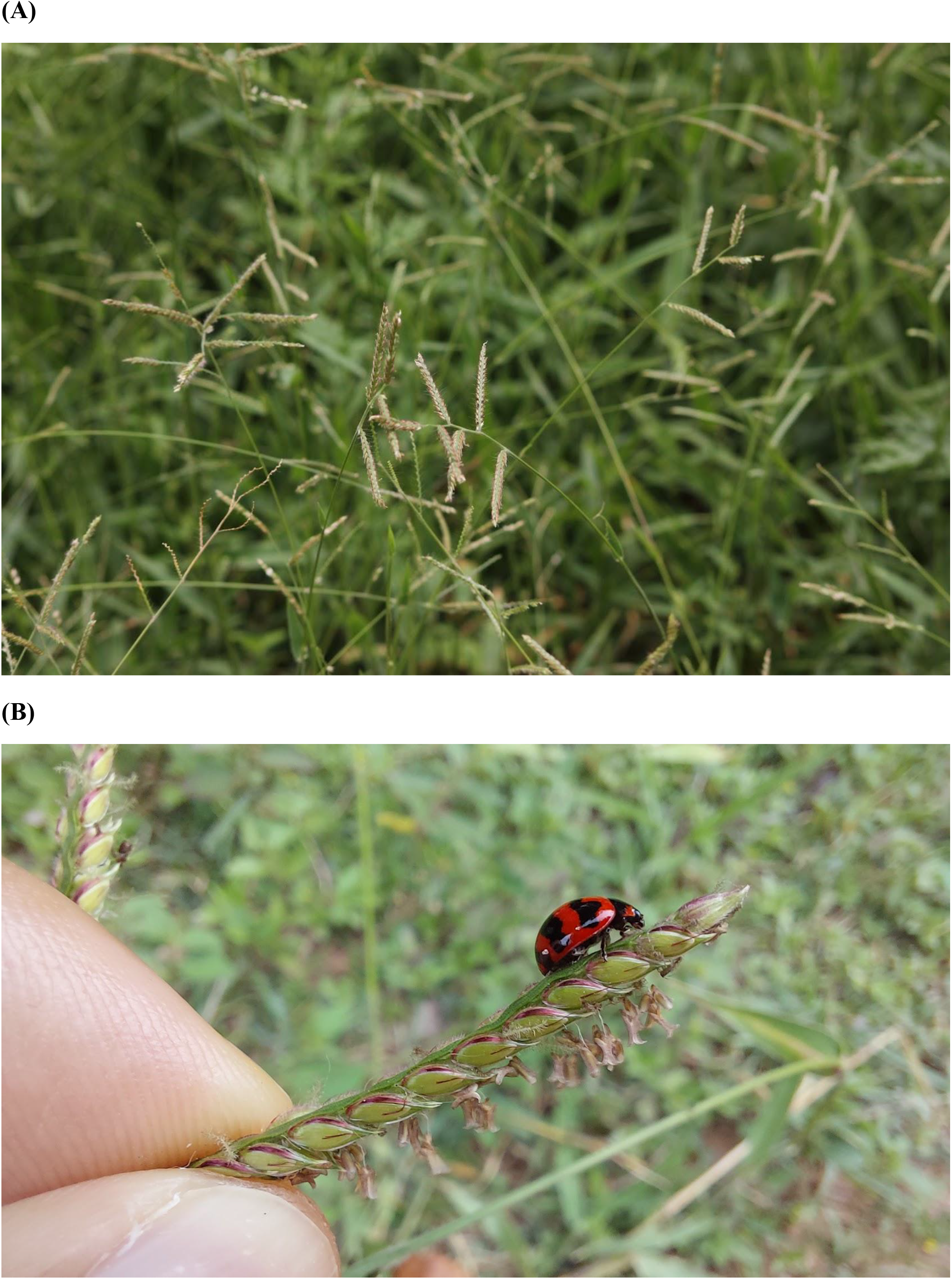
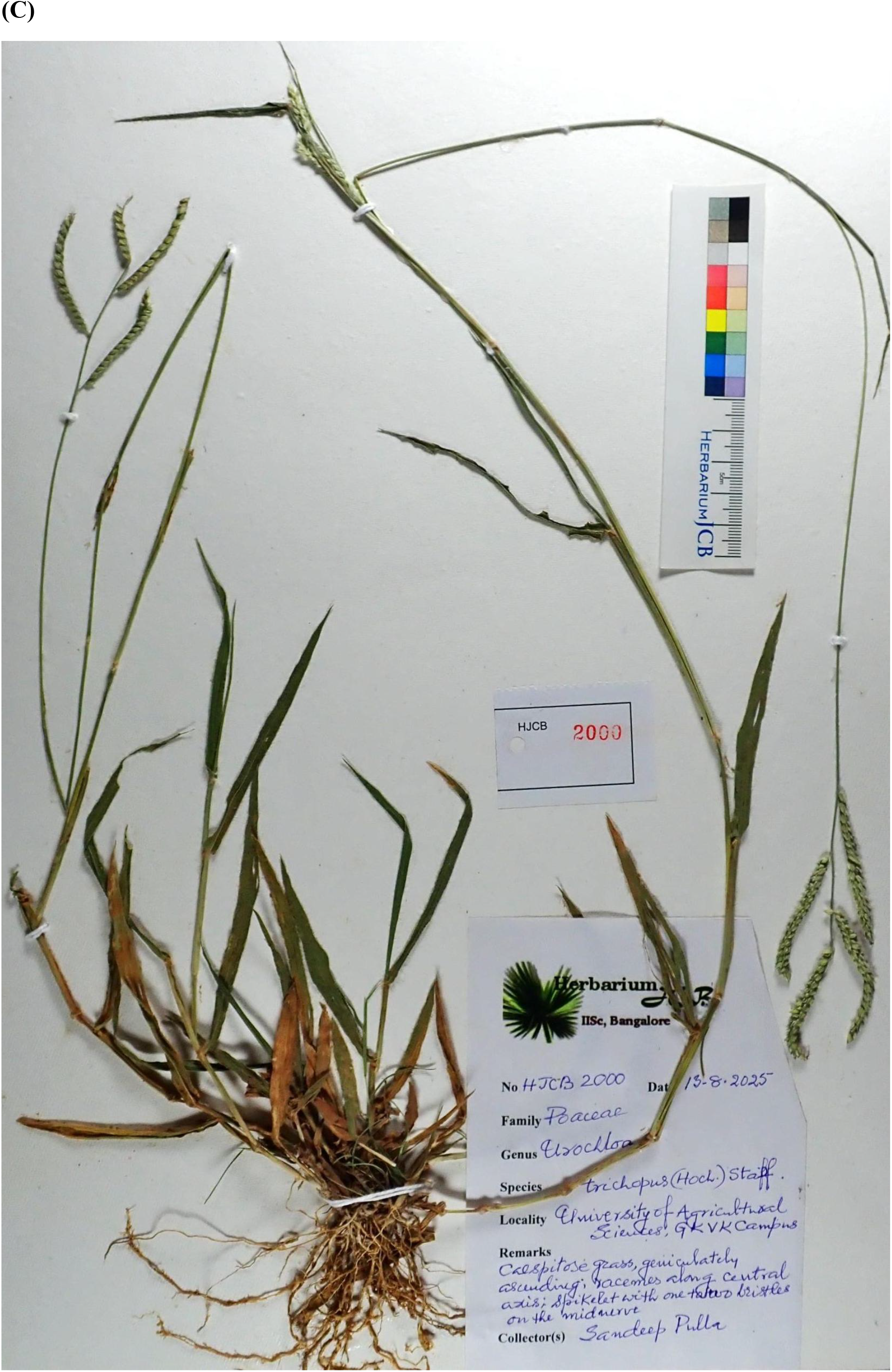
*Urochloa trichopus* (Hochst.) Stapf. (A) Inflorescences. (B) Raceme. (C) Herbarium specimen.

**Figure 2.**
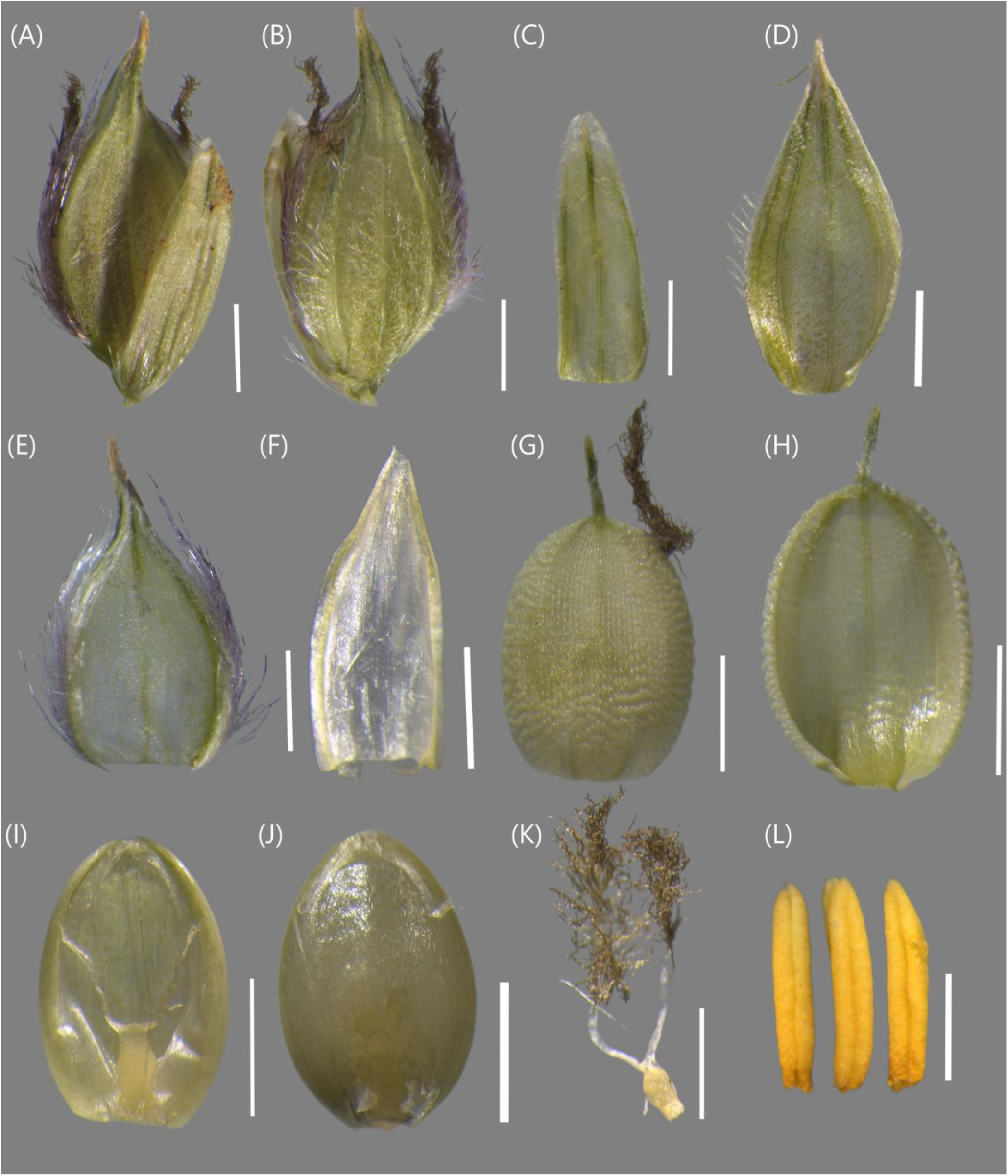
*Urochloa trichopus* (Hochst.) Stapf. (A) Spikelet abaxial view. (B) Spikelet adaxial view. (C) Lower glume. (D) Upper glume. (E) Lower lemma. (F) Lower palea. (G) Upper anthecium adaxial view showing germination flap in the lower-third. (H) Upper lemma abaxial view. (I) Upper palea with gynoecium. (J) Upper palea with caryopsis. (K) Gynoecium. (L) Anthers. All scale bars 1 mm.

## Notes

### Competing Interest Statement

The authors have declared no competing interest.

### Summary of Updates

Herbarium specimen image added and herbarium record cited; information on species ecology added.

## References

Ashalatha VN, Nair VJ (1993) Brachiaria Griseb. and Urochloa P.-Beauv. (Poaceae) in India - A conspectus. Nelumbo 27–31. 10.20324/nelumbo/v35/1993/74357

Bor NL (1960) The Grasses of Burma, Ceylon, India, and Pakistan, excluding Bambuseae. Oxford

Clayton WD, Renvoize SA (1986) Genera Graminum: Grasses of the world. H.M. Stationery Office

Delfini C, Salariato DL, Aliscioni SS, Zuloaga FO (2023) Systematics and phylogenetic placement of Panicum L. species within the Melinidinae based on morphological, anatomical, and molecular data (Poaceae, Panicoideae, Paniceae). Plants 12:399. 10.3390/plants12020399

Duque TS, da Silva RS, Maciel JC, Silva DV, Fernandes BCC, Júnior APB, Santos JB dos (2022) Potential distribution of and sensitivity analysis for Urochloa panicoides weed using modeling: an implication of invasion risk analysis for China and Europe. Plants 11:1761. 10.3390/plants11131761

Fish L, Mashau A, Moeaha M, Nembudani M (2015) Identification guide to southern African grasses: an identification manual with keys, descriptions and distributions. South African National Biodiversity Institute, Pretoria, South Africa

Gorgone-Barbosa E, Pivello VR, Bautista S, Zupo T, Rissi MN, Fidelis A (2015) How can an invasive grass affect fire behavior in a tropical savanna? A community and individual plant level approach. Biol Invasions 17:423–431. 10.1007/s10530-014-0740-z

Hartnett DC, Potgieter AF, Wilson GWT (2004) Fire effects on mycorrhizal symbiosis and root system architecture in southern African savanna grasses. Afr J Ecol 42:328–337. 10.1111/j.1365-2028.2004.00533.x

Heinl M, Sliva J, Murray-Hudson M, Tacheba B (2007) Post-fire succession on savanna habitats in the Okavango Delta wetland, Botswana. J Trop Ecol 23:705–713. 10.1017/S0266467407004452

Kabeer KAA, Nair VJ (2009) Flora of Tamil Nadu: Grasses. Botanical Survey of India Karnataka Biodiversity Board (2019) Flora of Karnataka, A checklist, Volume – 2:

Gymnosperms & Angiosperms. Karnataka Biodiversity Board, Karnataka

Kellogg EA (2015) Flowering Plants. Monocots: Poaceae. Springer

Kellogg EA, Abbott JR, Bawa KS, Gandhi KN, Kailash BR, Ganeshaiah KN, Shrestha UB, Raven P (2020) Checklist of the grasses of India. PhytoKeys 163:1–560. 10.3897/phytokeys.163.38393

Lawes RA, Grice AC (2010) War of the weeds: Competition hierarchies in invasive species. Austral Ecol 35:871–878. 10.1111/j.1442-9993.2009.02093.x

Masters LE, Tomaszewska P, Schwarzacher T, Hackel J, Zuntini AR, Heslop-Harrison P, Vorontsova MS (2024) Phylogenomic analysis reveals five independently evolved African forage grass clades in the genus Urochloa. Ann Bot 133:725–742. 10.1093/aob/mcae022

McIvor Joh NG, Howden SM (2000) Dormancy and germination characteristics of herbaceous species in the seasonally dry tropics of northern Australia. Austral Ecol 25:213–222. 10.1046/j.1442-9993.2000.01026.x

Pivello VR, Shida CN, Meirelles ST (1999) Alien grasses in Brazilian savannas: a threat to the biodiversity. Biodivers Conserv 8:1281–1294. 10.1023/A:1008933305857

Rojas-Sandoval J, Acevedo-Rodríguez P (2022) Megathyrsus maximus (Guinea grass). CABI Compend. 10.1079/cabicompendium.38666

Schindelin J, Arganda-Carreras I, Frise E, Kaynig V, Longair M, Pietzsch T, Preibisch S, Rueden C, Saalfeld S, Schmid B, Tinevez J-Y, White DJ, Hartenstein V, Eliceiri K, Tomancak P, Cardona A (2012) Fiji: an open-source platform for biological-image analysis. Nat Methods 9:676–682. 10.1038/nmeth.2019

Sharma BD (1984) Flora of Karnataka: Analysis. Botanical Survey of India, Department of Environment

Siddabathula N, Yadav PBS (2021) Urochloa trichopus Stapf, an addition to the flora of Telangana state, India

Sosef M (2016) Taxonomic novelties in Central African grasses (Poaceae), Paniceae 1. Plant Ecol Evol 149:356–365. 10.5091/plecevo.2016.1221

Treydte AC, Baumgartner S, Heitkönig IMA, Grant CC, Getz WM (2013) Herbaceous forage and selection patterns by ungulates across varying herbivore assemblages in a South African savanna. PLOS ONE 8:e82831. 10.1371/journal.pone.0082831

Van Oudtshoorn F (1999) Guide to grasses of southern Africa. Briza

Veldkamp J (1996) Brachiaria, Urochloa (Gramineae-Paniceae) in Malesia. Blumea Biodivers Evol Biogeogr Plants 41:413–437

Vorontsova MS (2022) Revision of some Malagasy forage grasses and their relatives within Brachiaria, Echinochloa, Moorochloa, and Urochloa. Candollea 77:199–236. 10.15553/c2022v772a7

Watson L, Macfarlane TD, Dallwitz MJ (1992) The grass genera of the world: descriptions, illustrations, identification, and information retrieval; including synonyms, morphology, anatomy, physiology, phytochemistry, cytology, classification, pathogens, world and local distribution, and references. Version: 15th June 2025. delta-intkey.com

Zenni RD, Ziller SR (2011) An overview of invasive plants in Brazil. Braz J Bot 34:431–446. 10.1590/S0100-84042011000300016

